# Regulation of global translation during the cell cycle

**DOI:** 10.1101/255729

**Authors:** Vilte Stonyte, Erik Boye, Beáta Grallert

## Abstract

It is generally accepted that global translation varies during the cell cycle and is low in mitosis. However, addressing this issue is challenging because it involves cell synchronization, which evokes stress responses which, in turn, affect translation rates. Here we have used two approaches to measure global translation rates in different cell-cycle phases. First, synchrony in different cell-cycle phases was obtained involving the same stress, by using temperature-sensitive mutants. Second, translation and DNA content were measured by flow cytometry in exponentially growing, single cells. We found no major variation in global translation rates through the cell cycle in either fission-yeast or mammalian cells. We also measured phosphorylation of eukaryotic initiation factor-2α, an event thought to downregulate global translation in mitosis. In contrast with the prevailing view, eIF2α phosphorylation correlated poorly with downregulation of general translation and ectopically induced eIF2α phosphorylation inhibited general translation only at high levels.

## Introduction

It is one of the basic principles of cell proliferation that there is a link between general cell growth (protein synthesis) and cell-cycle regulation. Such a link is logical and has been hypothesized to exist, but its nature has been elusive. Protein synthesis is one of the most energy-demanding cellular processes and is therefore carefully regulated. It is a generally accepted view that global translation is considerably reduced in mitosis (reviewed in 1). The reduction is thought to result from altered phosphorylation state of translation initiation factors. In particular, phosphorylation of the translation initiation factor eIF2α is induced after a number of different stresses and is thought to be the main reason for repressed translation. Cell-cycle-dependent downregulation of translation in G2/M phase was also attributed to increased eIF2α phosphorylation (2-5).

Early translation measurements in synchronized mammalian cells revealed a 70% reduction of the global translation rate in mitosis (6). More recent studies using different synchronization methods suggested that the magnitude of the translation reduction depends on the method of synchronization (7,8). Also, studies in budding yeast indicated that the rate of protein synthesis is constant during the cell cycle (9,10). More recent studies (in mammalian cells) have reported conflicting results regarding the level of translational reduction in mitosis (11-13), and the question of whether and to what extent global translation is downregulated in mitosis remains unanswered.

Measurement of translation in different cell-cycle phases is challenging because it often involves cell-cycle synchronization, which in itself can evoke stress responses which, in turn, will affect translation rates. Thus, the exact contribution of the synchronization method versus cell-cycle progression to any observed change in translation rates or the phosphorylation state of translation initiation factors is difficult to assess. Here we use novel approaches to measure global translation rates during the cell cycle and whether it depends on eIF2α phosphorylation.

## Results

### Global translation in synchronized cells

First, we utilized temperature-sensitive fission yeast mutants that arrest at different phases of the cell cycle. We synchronized the cells by shifting to the restrictive temperature before release into the cell cycle, achieving synchrony at different cell-cycle phases by the same treatment (ie temperature shift). Samples for analysis of DNA content, translation rate, and eIF2α phosphorylation were taken every 20 min for 160 or 220 minutes after release from the cell-cycle arrest. DNA content and translation rate were measured in single cells, by flow cytometry. Translation was assayed by pulse-labelling with the methionine analogue L-Homopropargylglycine (HPG) (15), which is incorporated into growing polypeptide chains. It should be noted that our assay addresses the regulation of global translation rather than the well-established translational regulation of individual proteins. To reveal small differences in signal intensity, the samples were barcoded and processed together in the very same solution. Phosphorylation of eIF2α was assessed by immunoblot analysis. The *cdc10-M17* mutant was used to synchronize cells in G1, *cdc25-22* was used to synchronize cells in G2 and *nda3-KM311* was used to arrest the cells in mitosis.

The rate of translation changed as the cells progressed from the block and through the cell cycle, apparently consistent with cell-cycle-dependent translation. However, the changes in translation rate followed the same pattern after release from the cell-cycle arrest regardless of when in the cell cycle the cells were synchronized (Fig 1A-D and Fig S1). At early time points the translation rate was low and after release it gradually increased to a rate above that measured before the shift. At late timepoints translation rates became similar to that measured in exponentially growing cells. There was no correlation between any particular cell-cycle phase and an increase or decrease in translation rates. These results strongly suggest that global translation is not regulated in a cell-cycle-dependent manner and that the variations observed are caused by the synchronization.

**Figure 1.**
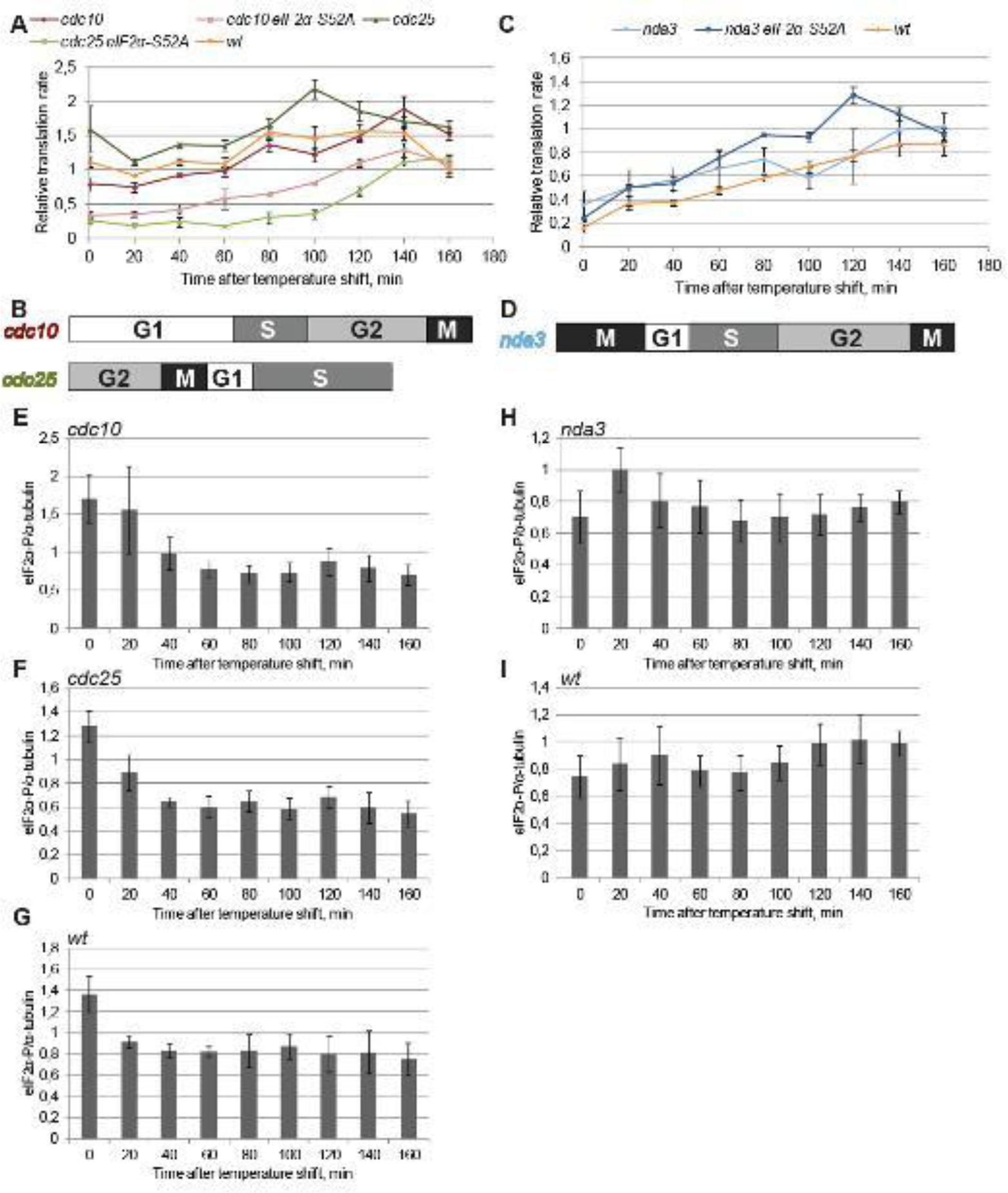
Global translation in cells synchronized in the cell cycle. Cells of the indicated strains were grown exponentially at 25°C **(A, B, E-G)** or 30 °C **(C, D, H, I)**, incubated at 36°C or 20°C for one generation time and then shifted back to 25 and 30 °C, respectively. Samples were taken at the indicated times after the shift. **A, C** median intensities of the AF647 (HGP) signal normalized to that of exponentially growing cells. Average of three biological repeats and standard errors (SE) are shown. **B, D** illustrate cell-cycle progression in the respective mutants. Fig S1 shows the cell-cycle distributions **E - I** Quantification of eIF2α phosphorylation normalized to tubulin in the indicated strains. Average and SE of three independent experiments are shown. Representative immunoblots are shown in Fig S2.

To test the effects of a temperature shift, wild-type fission yeast cells were subjected to the same shifts as employed to synchronize the cell-cycle mutants. Interestingly, translation rates followed the same pattern in the wild-type cells as in the cell-cycle mutants described above (Fig 1A, C), demonstrating that the observed changes are due to the temperature shift rather than to the cell-cycle stage where the particular mutant arrests. Furthermore, the temperature shift from 25 to 36 back to 25 °C in itself induced a transient G2 delay (Fig S1G), which is probably due to the previously described Rad3^ATR^-Rad9-dependent mechanism (14). Curiously, also a shift from 30 to 20 to 30°C induced a cell-cycle delay, but in G1/S (Fig S1H).

Phosphorylation of eIF2α was high at the early time points in the heat-sensitive mutants, then gradually diminished (Fig 1 E, F), regardless of where in the cell cycle the particular mutant was arrested. There was no correlation between eIF2α phosphorylation and any particular cell-cycle phase. As a control to assess synchrony achieved in the above experiments, we followed expression of the G1 cyclin Cig2 by immunoblotting. The previously reported cell-cycle-dependent regulation was obvious in all three strains (Fig S1), showing that the synchrony achieved in the above experiments allows us to detect cell-cycle-dependent changes in protein levels. Furthermore, the temperature shift resulted in increased eIF2α phosphorylation also in the wild-type cells (Fig 1G), confirming that such temperature shifts routinely employed in cell-cycle synchronization experiments invoke a stress response.

When cells were shifted from 20 to 30 oC, changes in eIF2α phosphorylation were much less pronounced, be it wild-type cells or the cold-sensitive *nda3* mutant (Fig 1H, I). Notably, the *nda3* mutant arrests in metaphase, the very cell-cycle phase where eIF2α phosphorylation is thought to increase and contribute to a downregulation of translation. Furthermore, the biggest change in translation rate was observed in the cells shifted from 20 to 30 oC, both for wild-type cells and the *nda3* mutant(Fig 1C), although this treatment resulted in the smallest change in eIF2α phosphorylation (Fig 1 H, I). These results are in direct contradiction to the prevailing view that eIF2α phosphorylation correlates with and is the reason for downregulation of global translation.

To assess the contribution of eIF2α phosphorylation to the observed changes in translation rates, strains carrying non-phosphorylatable *eIF2α-S52A* were used. Cell-cycle synchronization experiments and translation measurements were performed as above. Surprisingly, translation rates followed exactly the same pattern in the absence of eIF2α phosphorylation as in its presence; low immediately after the temperature shift, then recovering (Fig 1A, C). Furthermore, in the heat-sensitive mutants translation was much more downregulated when eIF2α could not be phosphorylated (Fig 1A).

We conclude that the changes in translation rates during the cell-cycle synchronization experiments were not due to cell-cycle-specific regulation of translation, but to the temperature shift itself. Furthermore, phosphorylation of eIF2α is not cell-cycle regulated and is not required for the downregulation of global translation after temperature shift.

### Global translation in exponentially growing cells

Having seen no evidence of cell-cycle dependent regulation of translation in synchronized cells, we set out to measure translation rates in different cell-cycle phases in unsynchronized cells. To this end, we measured HPG incorporation and DNA content in exponentially growing cells by flow cytometry. Cells in each cell-cycle phase were gated on two-parametric DNA cytograms (15) and HPG incorporation per cell was quantified in each cell-cycle phase. There were no significant differences in the rate of translation in the different cell-cycle phases (Fig 2A,C). It should be noted that this method does not allow us to distinguish cells in mitosis from those in G1. Thus, a high translation rate in G1 cells might compensate for a reduced translation rate in the mitotic cells so that the relative translation rate for the mixed M-G1 population appears to be unchanged. However, in such a scenario the distribution of the HPG intensities in the M-G1 population would be broad, but this is not the case (Fig 2A, C), arguing against this explanation. Another concern is that a low number of mitotic cells in the population would conceal a low translation rate in mitotic cells. To address this issue, cells of the M-G1 population were sorted onto microscopy slides and the microtubuli were stained. At least 20 % of the cells clearly contained a mitotic spindle (data not shown), demonstrating that the translation rates measured in the M-G1 population reliably represent those of mitotic cells. In addition, we analyzed exponentially growing fission yeast cells grown in a medium with isoleucine as sole nitrogen source. Under these conditions G1 is longer and cytokinesis occurs in G1 (16), which allows us to distinguish a G1 population containing 1C DNA from mitotic cells. Also under these conditions, translation rates were similar in the different cell-cycle phases (Fig 2B, D). These results obtained in unsyncronized, exponentially growing cells confirm that global translation does not vary significantly through the cell cycle.

**Figure 2.**
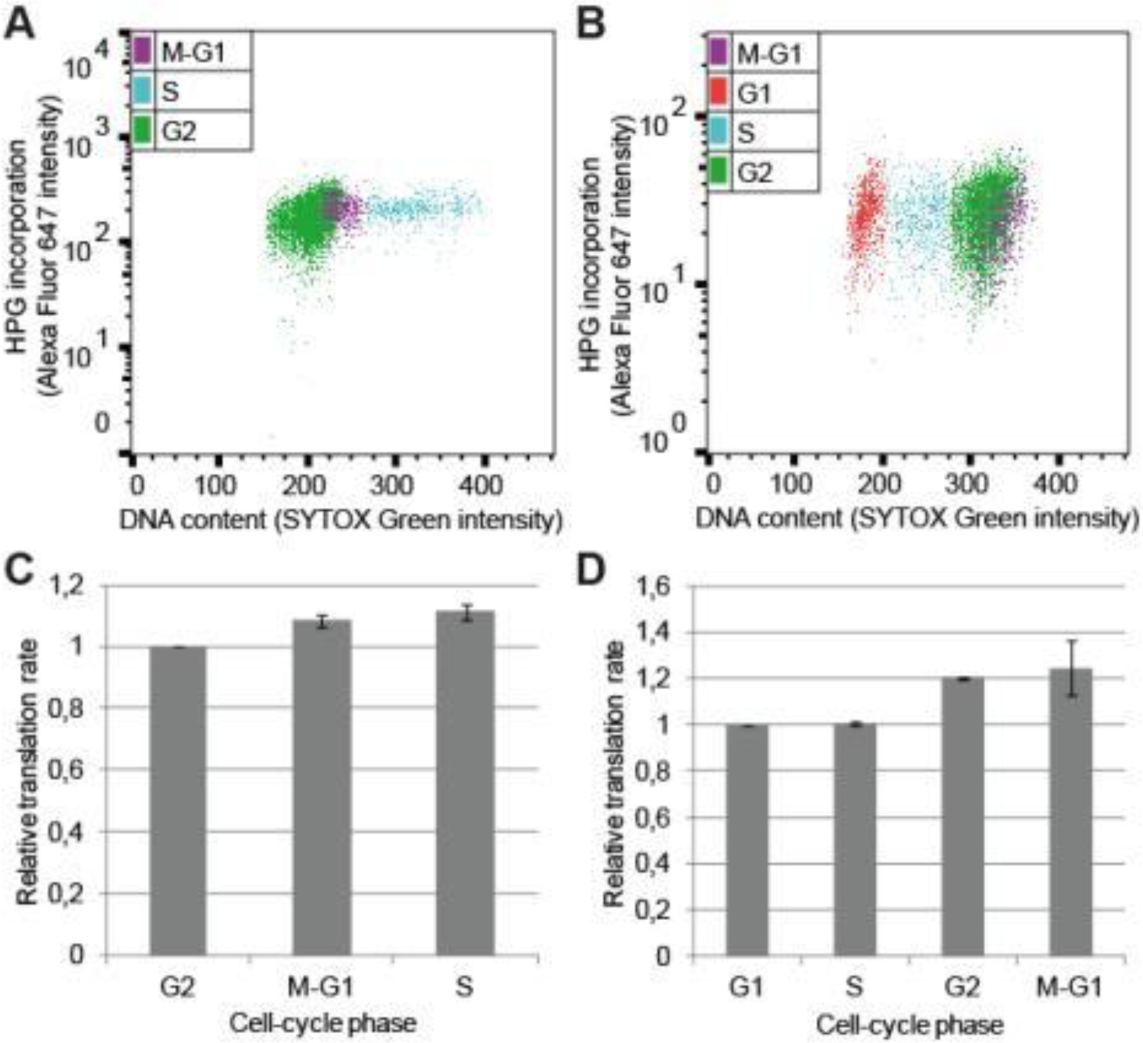
Global translation in exponentially growing cells. **A,B** Two-parametric flow cytometry plots of fission yeast cells grown in **(A)** EMM or **(B)** in isoleucine-minimal medium. **C, D** Average of median intensity of the AF647 signal normalized to G2 **(C)** or G1 **(D)** from at least three biological repeats with SE. Gating is shown on Fig S2.

Basic cellular processes such as regulation of translation through the cell cycle are expected to be conserved in evolution, but the extent of such regulation might vary from organism to organism. Therefore, we investigated whether the level of global translation varies during the cell cycle in human cells. To this end, we measured translation rates in different cell-cycle phases in three different human cell lines. To measure translation, unsynchronized cells were pulse-labelled with the puromycin analogue O-propargyl-puromycin (OPP) and analyzed by flow cytometry. Cells in G1, S and G2 were identified based on their DNA content and mitotic cells were identified using the mitotic marker phospho-S10-histone H3. The cell lines investigated were normal epithelial RPE cells immortalized by telomerase expression, the osteosarcoma-derived U2OS cells and cervix carcinoma-derived HeLa cells. There is a wide distribution of the intensity of the OPP signal in the G1 population, indicating that there are significant differences in translation rates among G1 cells. This feature is particularly obvious in the normal epithelial RPE cells, less pronounced in the two cancer cell lines (Fig 3). The G1 cells with lower translation rates might represent cells that have not yet passed the restriction point. There is a gradual increase in translation from G1 phase through S to G2 in all three cell lines, and a somewhat lower rate in mitotic cells. However, the rate of protein synthesis in mitotic cells is higher or similar to that in G1 cells and the extent of reduction from G2 to M ranges from 40% (RPE) to 15 % (U2OS).

**Figure 3.**
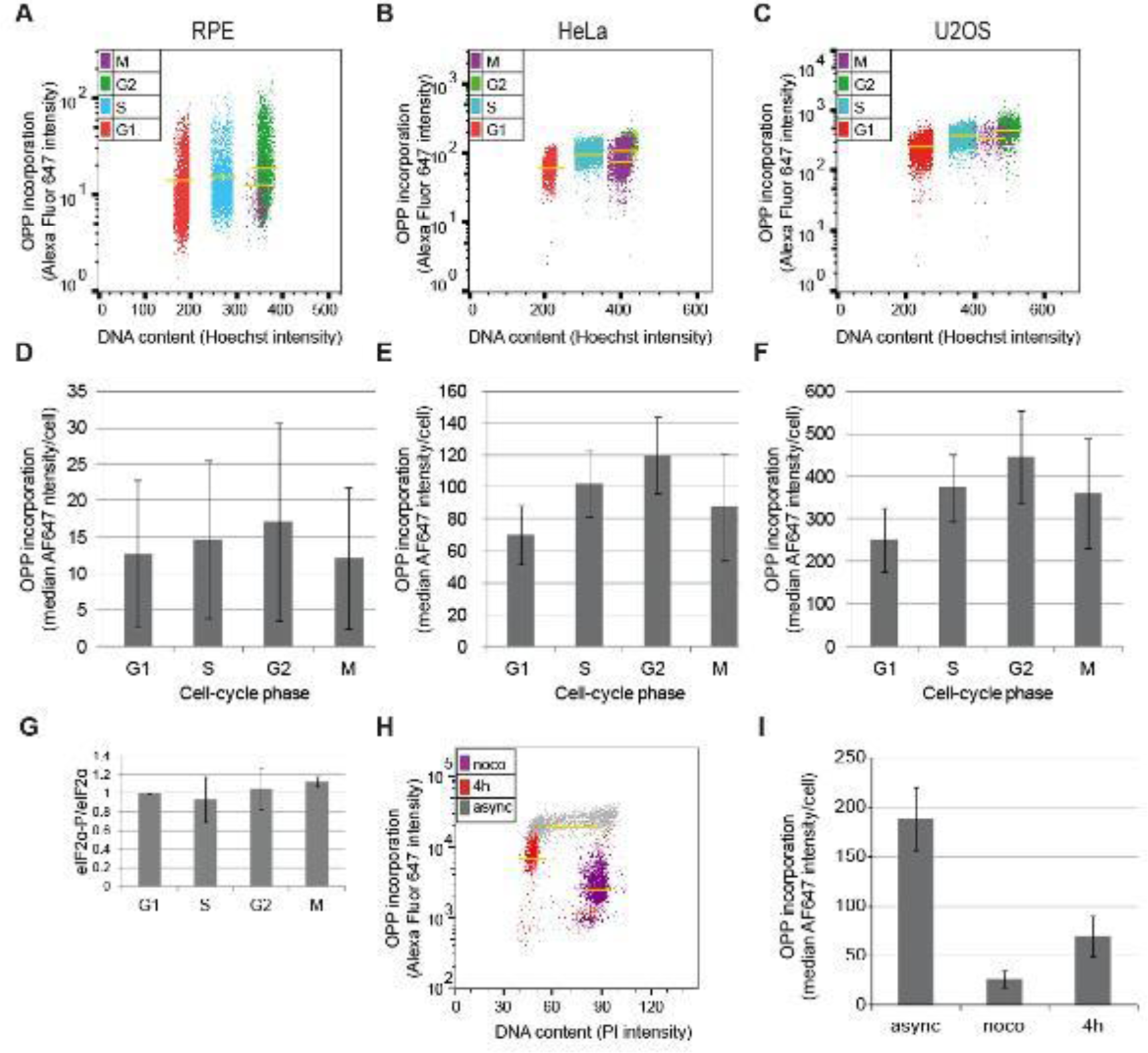
Global translation through the cell cycle in human cells. **A-C** Two-parametric flow cytometry plots of the indicated cell lines. Yellow lines represent the mean intensity of AF647 (OPP) for each cell-cycle phase. **D-F** Bar graphs representing mean AF647 (OPP) intensity with standard deviation. **G** Quantification of eIF2α phosphorylation normalized to eIF2α in the indicated cell-cycle phases. Exponentially growing HeLa cells were fixed and stained for H3-P and DNA content to identify cells in each cell-cycle phase and then 50 000 cells from each phase were sorted to measure eIF2α phosphorylation. Average and SE of three independent experiments are shown. Representative immunoblots are shown in Fig S3. **H.** Two-parametric flow cytometry plots of asynchronously growing and nocodazole-arrested cells and cells 4 h after release from the nocodazole block. **I** Bar graphs representing mean AF647 (OPP) intensity with standard deviation after nocodazole block and release. eIF2α phosphorylation is shown in Fig S3.

Phosphorylation of eIF2α was investigated in HeLa cells. Unsynchronized cells were fixed, analysed as above and collected by fluorescence-activated cell sorting (FACS). Phosphorylation of eIF2α was investigated in the different populations by immunoblotting. There were no significant changes in eIF2α phosphorylation during the cell cycle (Fig 3G).

The above results strongly suggest that the previously observed apparent cell-cycle-dependent variation in translation rates was a result of synchronization. In order to direcly address this, we synchronized HeLa cells using nocodazole and mitotic shake-off and measured the translation rates. Consistent with previous studies, translation rates changed dramatically in the nocodazole-tretated cells (Fig 3H, I) and eIF2α phosphorylation increased upon nocodazole arrest (Fig S3).

These findings strongly suggest that global translation rates are not dramatically downregulated in mitotic cells and that earlier studies overestimated the extent of variation through the cell cycle.

### eIF2α phosphorylation and general translation

Surprisingly poor correlation was observed between the levels of eIF2α phosphorylation and global translation in the temperature-shift experiments, prompting us to directly address the importance of eIF2α phosphorylation on global translation rates.

To this end, we expressed PKR, one of the four human eIF2α kinases, in fission yeast and measured eIF2α phosphorylation and the global translation rates. PKR expression was controlled by the regulatable *nmt1* promoter, which is induced when thiamine is removed from the medium (17,18). We used two different versions of the promoter, providing two different expression levels of PKR. Cells were grown exponentially with the promoter repressed before PKR expression was induced and global translation rates as well as eIF2α phosphorylation were measured during the first 24 hours (6 generations) after induction. PKR expressionwas detected at 13 hous after induction and eIF2α phosphorylation reached maximal values at 16 - 19 hours (Fig 4A, B and S4). The extent of eIF2α phosphorylation induced by PKR driven by the weaker promoter was comparable to that induced by milder stresses (Fig 4C and S4). Curiously, we did not see any significant decrease in global translation rates when PKR was expressed from the weaker of the promoters, the rate of translation remained similar to that before induction of PKR expression. (Fig 4D). However, in the cells expressing PKR from the full-strength *nmt* promoter translation was strongly reduced and, consistently, these cells could not form colonies when the promoter was derepressed (not shown). These results are consistent with previous findings, suggesting that extreme and lasting eIF2α phosphorylation can inhibit global translation and is lethal (19,20). We conclude that the extent of eIF2α phosphorylation is crucial for the effect on downregulation of general translation. A very high level of eIF2α phosphorylation blocks translation, but an intermediate level might have little influence on global translation.

**Figure 4.**
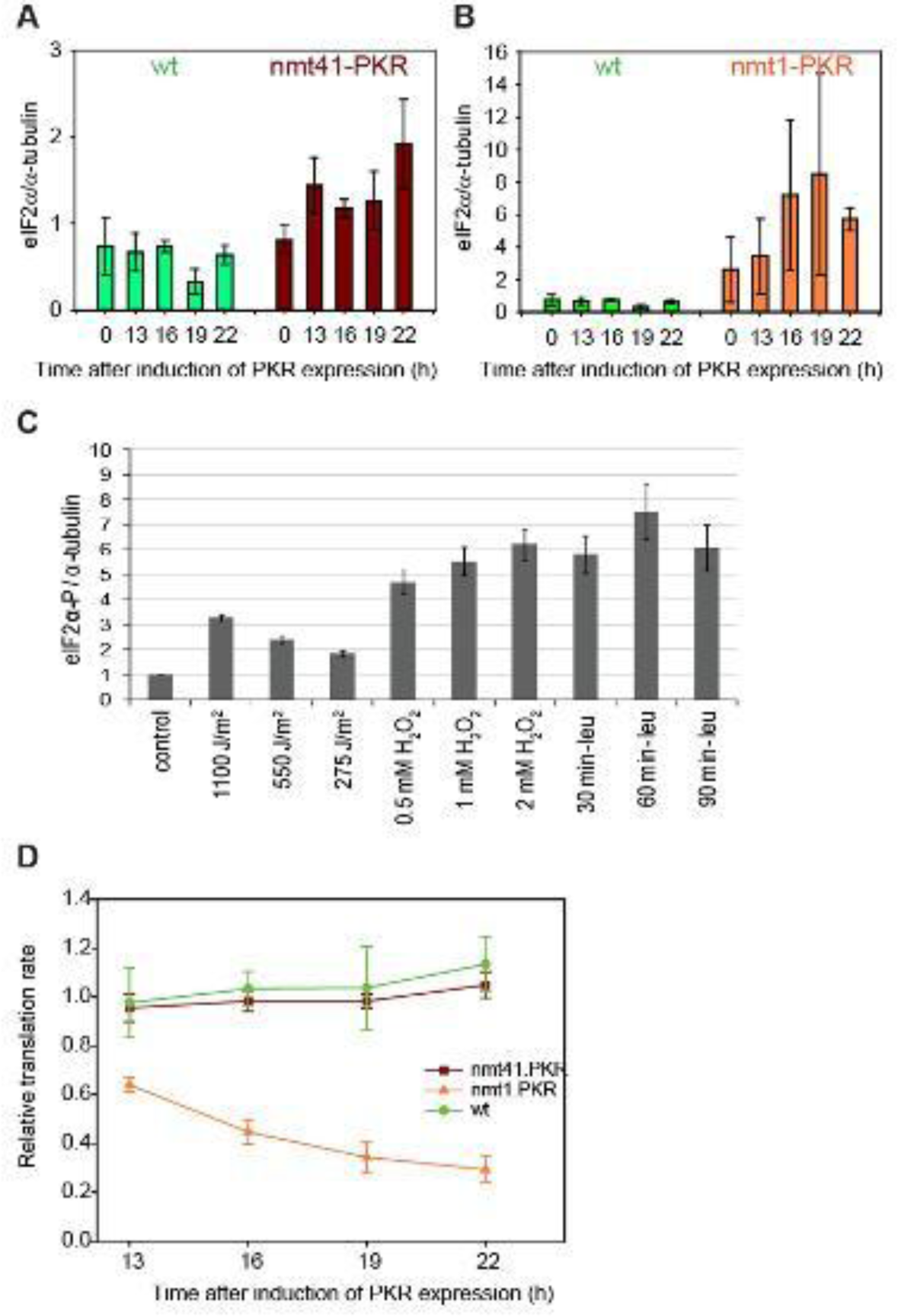
eIF2α phosphorylation and general translation. Cells carrying the indicated plasmids were grown exponentiallly with the promoter repressed and one sample was taken to measure translation. The promoter was induced for the indicated times. **A, B** Quantification of eIF2α phosphorylation normalized to α-tubulin at the indicated time points when PKR is expressed from the two different promoters. Note the different scales on the y axes. Representative immunoblots are shown in Fig S4. **C** Quantification of eIF2α phosphorylation normalized to tubulin after the indicated stresses. Average and SE of three independent experiments are shown. Representative immunoblots are shown in Fig S4. **D** Median intensities of the AF647 (HGP) signal normalized to that of exponentially growing cells (promoter repressed). Average of three biological repeats and SE are shown.

## Discussion

### Global translation rate changes little during the cell cycle

Many recent studies dispute the generally accepted view that global translation varies in a cell-cycle-dependent manner and is low in mitosis. Our results suggest that the discrepancies arise from experimental challenges. Studies of cell-cycle-related events often involve synchronization of cell cultures. In this work, we employed temperature-sensitive yeast mutants. It should be noted that studies on heat stress generally employ higher temperatures (>40 °C) and the temperatures we used are close to those in the natural environment of fission yeast cells. However, here we show that even the temperature shifts routinely used to synchronize the temperature-sensitive *S. pombe* mutants invoke a cellular stress response by themselves and influence global translation rates, supporting the idea that previously reported cell-cycle-dependent changes in translation rates are caused by the method of synchronization. Using the same stress to synchronize cells in different cell-cycle phases allowed us to separate the effects of cell-cycle progression from temperature shift on global translation rates. It is possible that, in our experiments, modest cell-cycle-dependent variations in global translation rates could be concealed by imperfect synchrony. However, the synchrony achieved in the block-and-release experiments (Fig S1) should have allowed us to observe the dramatic changes described previously. Furthermore, using flow cytometry to measure translation in exponentially growing cells allowed us to investigate global translation rates in different cell-cycle phases in unstressed cells.

One caveat of analyzing the cell cycle of fission yeast by flow cytometry is that mitotic cells can only be identified after separation of the daughter nuclei, but cells in the early phases of mitosis cannot be distinguished from cells in G2. Thus, a reduction of global translation rates in metaphase would not be detected using asynchronously growing cells and flow cytometry alone, although it would have been detected in the block-and-release experiments. Collectively, these data demonstrate that global translation is not significantly different between any of the cell-cycle phases in fission yeast cells.

In the human cell lines we also saw only small changes in the translation rate, consistent with recent studies reporting only minor variations. Mitotic cells were identified based on histone H3 phosphorylation, a mitotic marker that is present both in metaphase and anaphase. Notably, our approach did not involve any synchronization method, exposure to chemicals or changes in the cellular environment, which makes our results less subject to artifacts and methodical problems. Furthermore, when we synchronized the cells we also observed the previously reported variations, confirming the notion that the changes in translation are due to the synchronization-induced stress rather then cell-cycle progression.

### Phosphorylated eIF2α does not significantly repress global translation

Under stressful conditions cells reduce the rate of global translation to conserve resources (21). At the same time, synthesis of proteins necessary to survive the stress is maintained or even increased. Many different forms of stress results in phosphorylation of eIF2α in eukaryotic cells (22,23) and it is thought to be required for both responses; downregulation of general translation and upregulation of translation of selected mRNAs. In addition, it is also implicated in the cell-cycle-dependent regulation of translation. Here we find that increased eIF2α phosphorylation does not correlate with any particular cell-cycle phase, but rather with the stress involved in synchronization, be it temperature shift or exposure to nocodazole. We conclude that eIF2α phosphorylation is not regulated in a cell-cycle-dependent manner.

There is compelling evidence that eIF2α phosphorylation can attenuate the translation of mRNAs (24,25). The regulation of eIF2α phosphorylation is relevant for a number of diseases, such as neurodegenerative disorders, cancer and autoimmune diseases (26-31). In all these fields, increased levels of phosphorylated eIF2α has commonly been taken to be a readout of reduced general translation. However, the two parameters have rarely been measured in the same experiment. Our results demonstrate that there is poor correlation between eIF2α phosphorylation and repressed general translation. First, eIF2α phosphorylation is clearly not required for the temperature-shift-induced downregulation of translation (Fig 1), consistent with previous findings after UVC irradiation, oxidative stress and ER stress (32-34). Second, in the absence of eIF2α phosphorylation translation is repressed more dramatically after temperature shift (Fig 1). Third, ectopically induced eIF2α phosphorylation did not noticeably downregulate global translation in unstressed fission yeast cells, unless it was induced to high levels (Fig 4). We suggest that the impact of phosphorylated eIF2α on global translation has been overestimated in the literature and that eIF2α phosphorylation can not be used as a marker of downregulated translation. Our results demonstrate that the extent of eIF2α phosphorylation is crucial to determine whether it impacts on general translation and it has only a minor effect on the global translation at levels observed after mild stresses. This implies that the main consequence of eIF2α phosphorylation is not downregulation of general translation but most likely translation of selected mRNAs, as also suggested previously (35).

## Materials and Methods

### Cells and cell handling

All fission yeast strains used in this study are derivatives of S. pombe L972 h- wild-type strain (Leupold, 1950) and are listed in Table 1.

**Table 1.** Fission yeast strains used in this study.

|  |  |
| --- | --- |
| 19 | *L972 h-* |
| 489 | *cdc10-M17 h-* |
| 550 | *cdc25-22 h+* |
| 1711 | *nda3-KM311 cdt1:TAP:kanMX6 ura4-D18 h-* |
| 1244 | *cdc10-M17 eIF2alphaS52A:ura4+ ura4-D18 h+* |
| 2115 | *eIF2alphaS52A:ura4+ ura4-D18 nda3-KM311* |
| 2126 | *eIF2alphaS52A:ura4+ ura4-D18 cdc25-22* |
| 38 | *ura4-D18 leu1-32 h-* |

Cells were maintained and cultured as previously described (Moreno 1991). The cells were grown in liquid Edinburgh minimal medium (EMM) with appropriate supplements at 25 °C (or at 30 °C for *nda3-KM311*cells) to a cell concentration of 2-4× 10^6^/ml. The cells were synchronised in G1 or G2 phase by incubating *cdc10-M17* or *cdc25-22* cells, respectively, at 36 °C for 4 h (or 5 h for *cdc10-M17 eIF2alphaS52A* strain) before release into the cell cycle at 25 °C; in M phase by incubating *nda3-KM311* cells at 20 °C for 4 h before release into the cell cycle at 30 °C. To obtain a population of mononuclear G1 cells, cultures were maintained at 30 °C in minimal medium where NH_4_Cl was replaced with 20 mM L-isoleucine (Carlson et al., 1999). Cultures of *S. pombe* transformants (together with a wild-type control culture) were grown to a cell concentration of 8× 10^6^/ml (OD_595_ = 0.4) in minimal medium where NH_4_Cl was replaced with 3.75 g/l L-glutamic acid, monosodium salt (Pombe Glutamate medium, PMG). To induce human PKR expression, cells cultured in PMG containing 5 µg/ml thiamine (Sigma-Aldrich) were harvested by centrifuging for 3 min at 3 000 rpm, washed three times with PMG without thiamine, and resuspended in PMG lacking thiamine for the induction of *nmt1* and *nmt41* promoters.

Human HeLa and U2OS cells were cultivated in Dulbecco’s modified Eagle’s Medium (DMEM) (Gibco) and Tert-RPE cells were cultivated in DMEM-F-12 (Gibco) supplemented with 10% fetal bovine serum and 1% Penicillin/Streptomycin at 37°C in a humidified environment with 5% CO2.

### Cell-cycle analyses

Cell-cycle phases were identified in fission yeast by DNA staining (Sytox Green) as described (15). In mammalian cells DNA staining (Hoechst or propidium iodie) and phospho-histone H3 (Ser10) staining were employed.

### Translation assays

To label newly synthesized proteins, 50 µM of L-homopropargylglycine (HPG, Thermo Fisher Scientific) was added to 1 ml samples of the main yeast culture taken out 10 min before the indicated time points. To stop translation, 0.1 mg/ml of cycloheximide (CHX) was added after 10 min. Cells were fixed in ice-cold methanol or 70 % ethanol, washed in 0.5 ml TBS and barcoded using up to five different concentration (450, 124.8, 31.2, 6.24, and 0.78 ng/ml) of Pacific Blue (PB; Thermo Fisher Scientific) dye for 30 min in the dark at room temperature. Samples were then washed three times in 0.5 ml TBS and pooled together. The samples were permeabilised with 0.5 ml 1 % Triton X-100 in TBS, and blocked with 1 % BSA in TBS. To detect HPG, Alexa Fluor 647 was linked to the incorporated HPG in a ‘click’ reaction (Liang, Astruc, 2011) using the Click-iT cell reaction buffer kit (Thermo Fisher Scientific C10269) following the manufacturer’s protocol to ligate the HPG alkyne with a fluorescent azide. Incorporation was quantified by using flow cytometry (LSR II flow cytometer, BD Biosciences). SYTOX Green dye (Thermo Fisher Scientific) was used to stain the DNA. Cell doublets were excluded from the analysis as described previously (Knutsen, 2011). Samples without HPG were used as negative controls.

O-propargyl-puromycin (OPP), (Thermo Fisher Scientific) was added to 6 µM for 20 min, the cells were then trypsinized and fixed in 70% ethanol. To detect incorporated OPP, the fixed cells were washed once in PBS with 1 % FBS. OPP was ligated with Alexa Fluor 647 in a ‘click’ reaction following the manufacturer’s instructions. The samples were incubated for 5 min in detergent buffer (0.1 % Igepal CA-630, 6.5 mM Na_2_HPO_4_, 1.5 mM KH_2_PO_4_, 2.7 mM KCl, 137 mM NaCl, 0.5 mM EDTA (pH 7.5)) containing 4 % non-fat milk to block non-specific binding. The cells were incubated for 1 h with anti-phospho-histone H3 (Ser10) primary antibody (1:500, Millipore 06-570) in detergent buffer containing 2 % non-fat milk, washed once in PBS with 1 % FBS, and incubated for 30 min with Alexa Flour 488-linked secondary antibody (1:500, Thermo Fisher Scientific A-11034) in detergent buffer. All incubations were carried out in the dark at room temperature. The cells were washed once in PBS with 1 % FBS and stained with 1.5 μg/ml of Hoechst 33258 (Sigma) in PBS. The samples were analysed using flow cytometry (LSR II flow cytometer, BD Bioscience, San Jose, CA, USA). Samples without OPP and without the primary antibody were used as negative controls.

### Fluorescence activated cell sorting

Exponentially growing cells were fixed with 70% EtOH and stained for anti-phospho-histone H3 as described above and detected using Alexa-fluor 647-coupled secondary antibody. DNA was stained with 8 μg/ml propidium iodide. 50 000 cells from each cell-cycle phase were harvested using a FACS Aria II cell sorter.

### UVC irradiation

Fission yeast cells were irradiated with 254 nm UV light (UVC) in a suspension in EMM (or PMG) medium under continuous stirring to ensure equal irradiation dose (Nilssen, 2003). The incident dose was measured with a radiometer (UV Products). A surface dose of 1100 J/m^2^ (at a dose rate of approximately 250 J/m^2^/min) induces a checkpoint response, but results in over 90 % cell survival. Samples for protein analysis were taken immediately after irradiation.

### H_2_O_2_ treatment

Cells grown in PMG medium were treated with H_2_O_2_ at the indicated concentrations for 15 minutes before samples were taken.

### Leucine starvation

An auxotroph strain was grown in PMG medium supplemented with leucine. The cells were washed with PMG medium three times and incubated in medium not containing leucine for the indicated times.

### Immunoblotting

Total protein extracts of yeast cells were obtained using a low salt buffer (25 mM MOPS (pH 7.1), 60 mM β-glycerophosphate, 15 mM p-nitrophenylphosphate, 15 mM MgCl_2_, 15 mM EGTA (pH 8.0), 1 mM DTT, 0.1 mM Na_3_VO_4_, 1 % Triton X-100) supplemented with protease inhibitors (Roche). Cell debris was removed by centrifugation at 14 000 g for 15 min at 4 °C. The extracts were mixed with 4× LDS Sample Buffer (Thermo Fisher Scientific) and 50 mM DTT. Human cells were lysed in Laemmli sample buffer. Extracts were run on polyacrylamide gels, transferred onto PVDF membranes, and probed with antibodies against phospho-eIF2α (1:750, CST 3398), eIF2α (1:1000, Santa Cruz sc-11386) PKR (1:3000, Abcam 32052), α-tubulin (1:30 000, Sigma-Aldrich T5168) and γ-tubulin (1:30 000,). The signal intensities were quantified using ImageJ software.

## Acknowledgements

We are grateful to the Norwegian Cancer Society, the Norwegian South-Eastern Health Authority and Radiumhospitalets Legater for funding and thank L. Lindbergsengen and M. O. Haugli for excellent technical assistance. The funders had no role in study design, data collection and interpretation, or the decision to submit the work for publication.

## Supplementary figure legends

**Fig S1, related to Fig 1.**
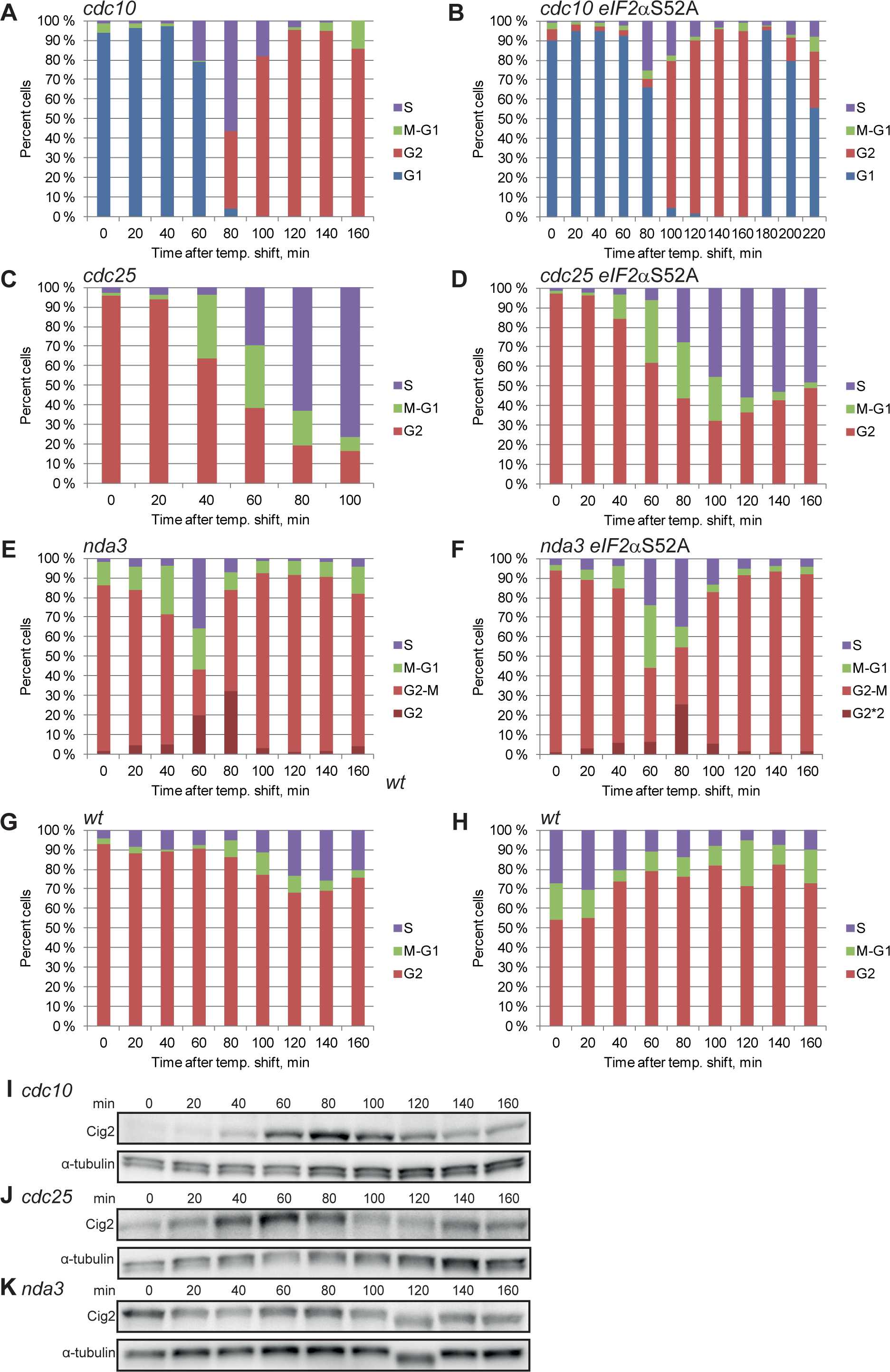
**A-H** Cell-cycle progression in the cell-cycle synchronization experiments. **I-K** Representative immunoblots of the cell-cycle-regulated Cig2 cyclin in the indicated mutants. α-tubulin is shown as loading control.

**Fig S2, related to Fig 1 and Fig2.**
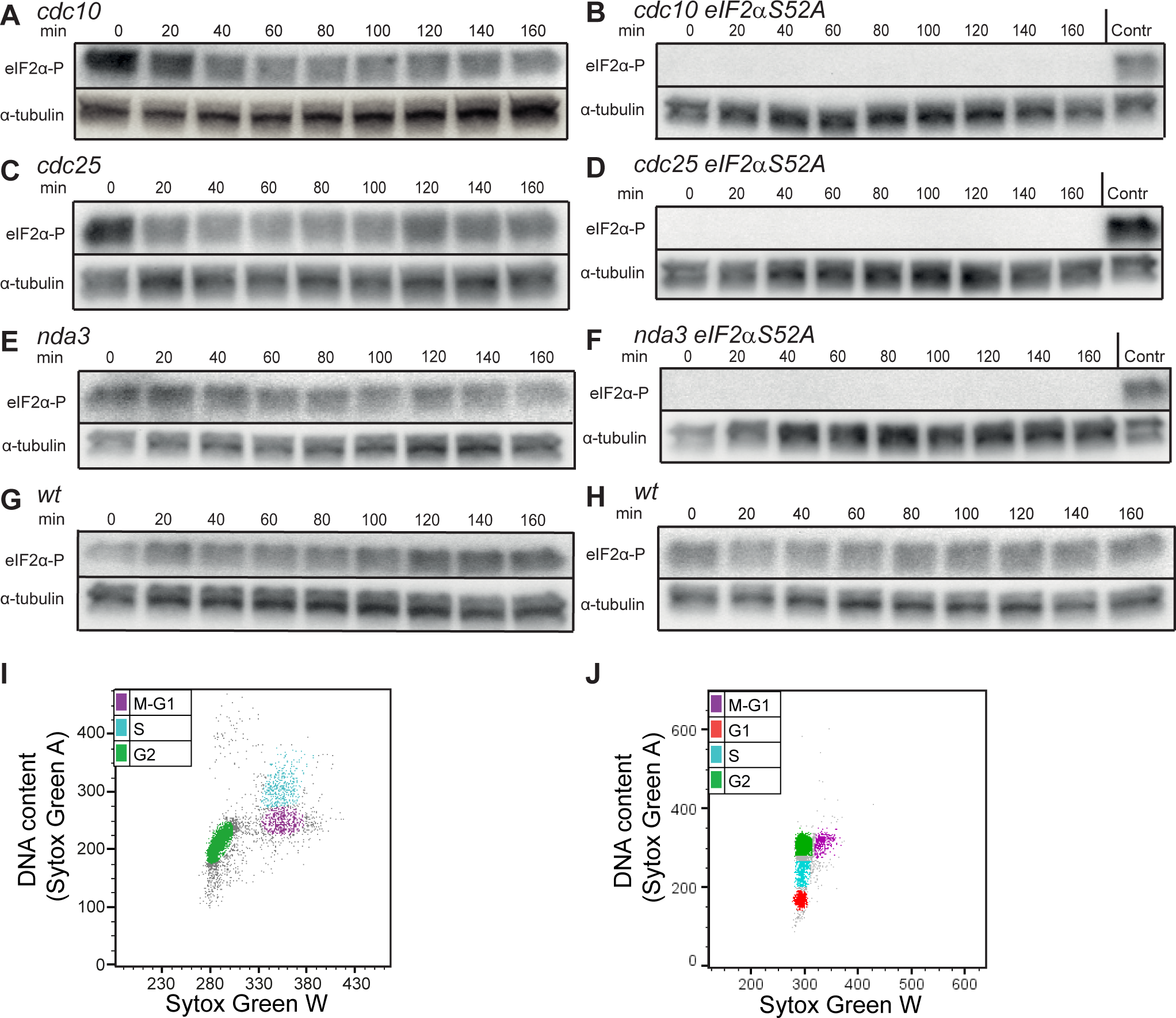
**A - H** Representative immunoblots showing eIF2α phosphorylation and tubulin loading control in the indicated strains. **I-J** Two-parametric DNA histograms showing gating used for the plot shown in Fig. 2A, B.

**Fig S3 related to Fig. 3.**
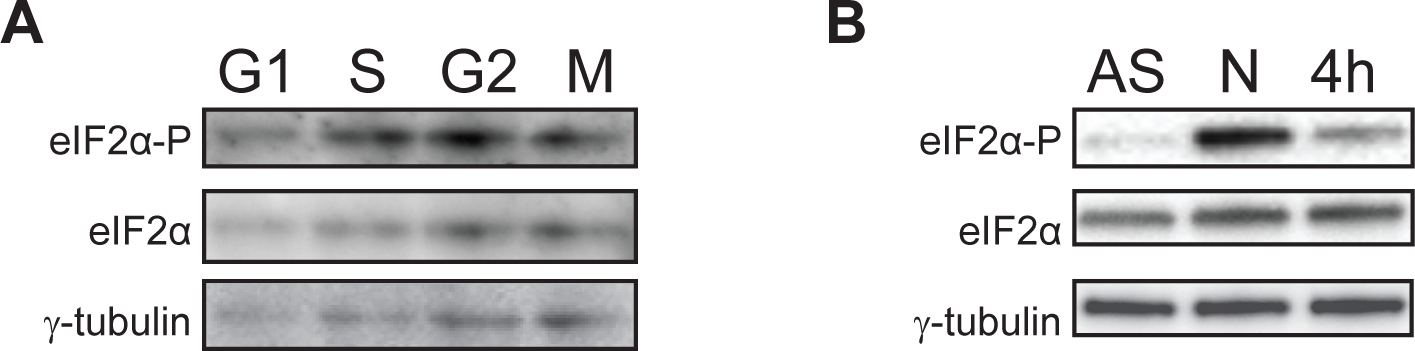
**A** eIF2α phosphorylation in the different cell-cycle phases. **B** eIF2α phosphorylation in asynchronous (AS) and nocodazole-arrested cells (N) and 4h after release from a nocodazole arrest (4h).

**Fig S4 related to Fig 4.**
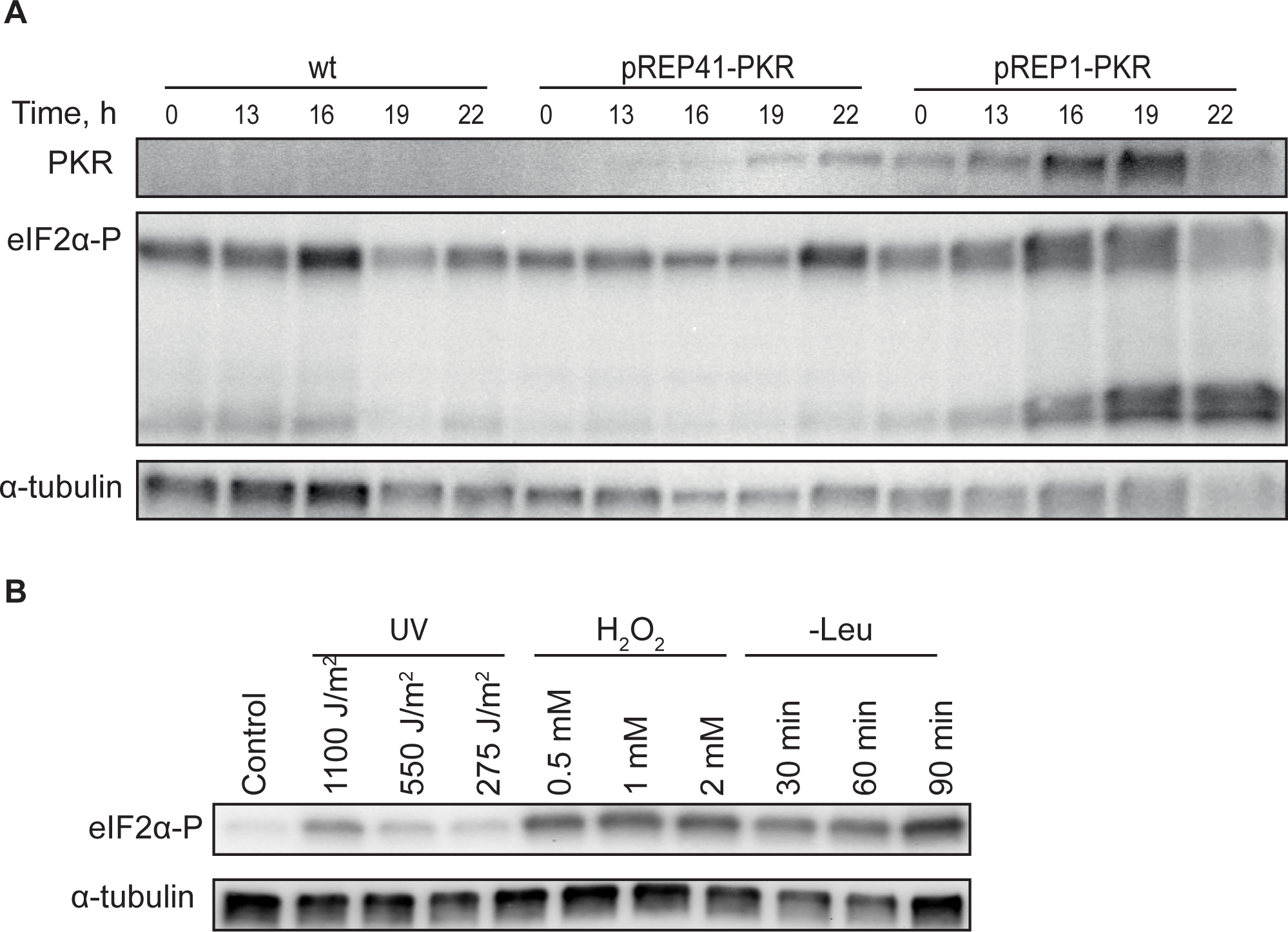
**A** Representative immunoblots showing PKR expression, eIF2α phosphorylation and loading controls in the indicated strains. **B** Representative immunoblot showing eIF2α phosphorylation and loading control in a wild-type strain after treatment with the indicated stresses.

